# *Npas4a* expression in the teleost forebrain is associated with stress coping style differences in fear learning

**DOI:** 10.1101/2020.11.24.396887

**Authors:** Matthew R Baker, Ryan Y Wong

**Author notes:** Correspondence: Ryan Y Wong, University of Nebraska at Omaha, 6001 Dodge St, Omaha, NE 68182. **Significance Statement:** Learning to predict and cope with potentially dangerous environments is an adaptive survival response. Proactive and reactive stress coping styles represent alternative strategies for coping with stress and differ in a number of behavioral contexts, including learning and memory. We show that reactive zebrafish display stronger conditioned fear responses to an olfactory alarm cue, with associated higher expression of a neuroplasticity-related gene, *npas4a*, in the medial and lateral zones of the dorsal telencephalon, and the supracommissural nucleus of the ventral telencephalon. Our study suggests that *npas4a*-dependent plasticity in the teleost forebrain is important for individual variation in fear learning. More broadly, plasticity in these associative limbic regions may regulate alternative stress coping styles and constrain behavioral variation across a number of behavioral contexts.

## Abstract

Learning to anticipate potentially dangerous contexts is an adaptive behavioral response to coping with stressors. An animal’s stress coping style (e.g. proactive-reactive axis) is known to influence how it encodes salient events. However, the neural and molecular mechanisms underlying these stress coping style differences in learning are unknown. Further, while a number of neuroplasticity-related genes have been associated with alternative stress coping styles, it is unclear if these genes may bias the development of conditioned behavioral responses to stressful stimuli, and if so, which brain regions are involved. Here, we trained adult zebrafish to associate a naturally aversive olfactory cue with a given context. Next, we investigated if expression of two neural plasticity and neurotransmission-related genes (*npas4a* and *gabbr1a*) were associated with the contextual fear conditioning differences between proactive and reactive stress coping styles. Reactive zebrafish developed a stronger conditioned fear response and showed significantly higher *npas4a* expression in the medial and lateral zones of the dorsal telencephalon (Dm, Dl), and the supracommissural nucleus of the ventral telencephalon (Vs). Our findings suggest that the magnitude of expression of activity-dependent genes like *npas4a* may be differentially expressed across several interconnected forebrain regions in response to fearful stimuli and promote biases in fear learning among different stress coping styles.

## Introduction

Animals frequently must overcome stressors and the ability to encode and recall these salient experiences is essential to an individual’s survival. Within individuals, behavioral and physiological responses to stressors often co-vary, belonging to correlated suites of traits that are consistent across contexts and time(1–4) (i.e. animal personality, stress coping styles; bold-shy axis, proactive-reactive axis). In addition to boldness, aggression, and stress physiology, studies demonstrate that proactive and reactive individuals also differ in learning and memory processes(5–9). The more risk-prone proactive individuals tend to show faster acquisition of memories that require higher levels of activity, or paradigms with positive and rewarding valence(10–16). In contrast, the risk-averse reactive individuals tend to show faster acquisition of aversive paradigms that require avoidance or reduced levels of activity(17–19). Despite these findings, the neuromolecular mechanisms and regional brain activity underlying these stress coping style differences in learning are not well understood.

Recent work has suggested that neural plasticity and neurogenesis may be key mechanisms underlying divergent proactive-reactive responses to stress, but whether these processes are associated with differences in learning and memory is not understood (20, 21). While previous studies have characterized the whole-brain transcriptome of proactive and reactive individuals at baseline, the contribution of specific neural plasticity- and synaptic transmission-related candidate genes and their spatial expression patterns have yet to be examined during a learning and memory task (22, 23). Two particularly interesting candidate genes, *npas4* and *gabbr1 (npas4a* and *gabbr1a* in teleosts) are essential in regulating neuronal excitability and molecular processes related to learning and memory such as long-term potentiation (24–26). *npas4* is an immediate early gene transcription factor that is predominantly expressed in the brain and enriched in the limbic regions. It is expressed through calcium signaling and is thought to induce primarily GABAergic inhibitory synapses in response to excitation and play an important role in homeostatic plasticity(25). *gabbr1* codes for a metabotropic GABA B receptor, which has also been shown to play an important role reducing neuronal excitability through G-protein signaling-dependent slow, long lasting hyperpolarization of postsynaptic cells. Further, deletion or altered expression of both of these genes has been shown to cause abnormal synaptic plasticity, neurogenesis, and impaired learning and memory abilities(27–29). Both of these genes were found to have significantly upregulated whole-brain expression at baseline in selectively-bred reactive zebrafish, which separately showed faster acquisition of a contextual conditioned fear response towards an aversive olfactory alarm cue (alarm substance)(22, 30). However, it is unknown if expression of these genes in specific brain regions are more directly associated with proactive-reactive differences in fear learning.

The basic neural substrates of fear learning have been well characterized, and are promising candidate sites where neural plasticity-related processes may regulate variation in fear learning capabilities. Traditionally, the basolateral amygdala is at the center of the fear system, with the hippocampus providing relevant associative information to allow for context-specific defensive responses fearful stimuli(31). More recently other brain regions such as the bed nucleus of the stria terminalis (BNST), lateral septum (LS), and striatum have attracted greater interest due to their functional and structural connections with the hippocampal/amygdala affective forebrain, and their output to structures essential for behavioral and physiological responses to potential threats. The majority of this circuitry has been characterized in rodent models, with putatively homologous structures identified in the teleost forebrain which have also been shown to be critical for contextual fear learning and adaptive responses to stress(32–35).

Here, we trained proactive and reactive zebrafish to associate alarm substance exposure with a context in one training trial, followed by a second assessment trial in the absence of the alarm substance. We then quantified *npas4a* and *gabbr1a* forebrain expression to investigate their potential link with differences in conditioned fear responses between alternative stress coping styles. We predict that an increased conditioned fear response in reactive zebrafish will be associated with increased expression of neural plasticity-related genes in the dorsal and medial portions of the dorsal telencephalon (Dm, Dl) and the dorsal, ventral, and supracomissural portions of the ventral telencephalon (Vd, Vv, Vs), putative homologues of the mammalian basolateral amygdala, hippocampus, striatum, lateral septum, and bed nucleus of the stria terminalis, respectively(32–35).

## Methods

### Subjects

Zebrafish are utilized in a variety of laboratory studies to understand the neural, genetic, and pharmacological mechanisms of learning and memory(36–38). Both wild and laboratory strains of zebrafish display the proactive and reactive stress coping styles, which have distinct genetic architectures and neuroendocrine responses (22, 23, 39). Here we used the high-stationary behavior (HSB; reactive) and low-stationary behavior (LSB; proactive) zebrafish strains to study the association between *npas4a* and *gabbr1a* expression and fear learning differences between proactive and reactive stress coping styles. Starting from wild-caught zebrafish, the HSB and LSB strains were generated and are maintained by artificial selection for opposing amounts of stationary behavior to a novelty stressor(40). The HSB and LSB strains show contrasting behavior, physiology, morphology, and neuromolecular profiles consistent with the reactive and proactive coping styles, respectively(22, 40–44). Additionally, these divergent behavioral profiles between the strains are consistent across contexts and over time and have high repeatability (40, 45, 46). During testing, fish were individually housed in 3-liter tanks on a recirculating water system (Pentair Aquatic Eco-Systems) using UV and solid filtration on a 14:10 L/D cycle at a temperature of 27° C. Fish were fed twice a day with Tetramin Tropical Flakes (Tetra, USA).

### Alarm Substance

We created a single batch of alarm substance as previously described(30). In brief, 20 randomly selected donor fish (wild type) were euthanized by rapid chilling followed by light abrasion of lateral skin cells on one side of each donor fish, ensuring that no blood was drawn. Donor bodies were then individually soaked in 10 mL of DI water for 10 minutes. A total of 200 mL was filtered, diluted in half, and stored in aliquots at −20° C until use. All procedures were approved by the Institutional Animal Care and Use Committee of University of Nebraska at Omaha/University of Nebraska Medical Center (17-070-00-FC, 17-064-08-FC).

### Contextual Fear Learning

To test learning, we utilized a validated contextual fear conditioning paradigm (30). Briefly, zebrafish were tested individually in a 16 × 16 × 10 cm arena filled with 1.4 L of system water. The arena was surrounded by opaque white plastic on the bottom and sides to serve as the contextual stimulus. Animals were removed from group housing and placed into individual housing 72 hours prior to the training session. Each learning trial was 15 minutes long and was divided into three subsections. Fish acclimated to the chamber for the first five minutes, followed by five minutes of recording pre-exposure behavior (conditioned fear response for second trial). After these 10 minutes, 1 mL of alarm substance (AS) or distilled water (DI) was administered into the water through plastic tubing that came from outside of the testing arena. Following alarm substance exposure, the unconditioned fear response was recorded for five minutes. Between trials, fish were placed back into their individual housing, the testing arenas were rinsed out, and were refilled with 1.4 L of fresh system water. Fish underwent two training trials with 30 minutes between trials. The second training trial was stopped after the second five minute block (conditioned response). Fish immediately had their forebrains removed or were decapitated and frozen on dry ice and stored at −80°C for qPCR and ISH, respectively. We selected the second trial for gene expression analyses because we previously showed that out of four training trials, the second trial was both the earliest trial and one that resulted in the most prominent proactive-reactive behavioral differences during fear conditioning before both lines achieved similar conditioned responses. These differences during training were also associated with stronger fear memory recall 96h following training (30).

Total sample sizes consisted of 46 LSB (N = 28 males, 18 females) and 46 HSB (N = 28 males, 18 females) individuals. Of this total, we used 10 HSB individuals (N = 5 AS, 5 DI, all males) and 10 LSB individuals (N = 5 AS, 5 DI, all males) for qRT-PCR analysis. We used the remaining fish for ISH analysis. A total of 12 LSB (N = 6 males, 6 females) and 12 HSB (N = 6 males, 6 females) individuals received alarm substance CS-US reinforcements as the experimental group. For the DI water control group, we used 12 HSB (N = 6 males, 6 females) and 12 LSB (N = 6 males, 6 females) fish. To control for possible effects of the paradigm and handling, independent of treatment group, 12 HSB (N = 6 males, 6 females) and 12 LSB (N = 6 males, 6 females) were habituated to the same single housing as other groups, but did not undergo behavioral testing.

### Behavioral Analysis

All trials were video-recorded from above and later analyzed with Noldus Ethovision XT (Noldus XT, Wageningen, Netherlands). For each trial, we quantified freezing time as an indicator of the conditioned response. We examined freezing because it is one of the most consistent and conserved behaviors used to assess stress-related behaviors and fear learning and memory(47). Additionally, freezing was the most reliable indicator of proactive-reactive differences in contextual fear conditioning in our prior study(30). The subject was considered frozen if it moved less than 0.5 cm/s.

### qRT-PCR

Preparation, execution, and analysis of the qRT-PCR of forebrain *npas4a* and *gabbr1a* expression followed previously established methods(42, 43). Gene expression was normalized to an endogenous housekeeping gene, *ef1a*, which has shown to be stable across sex, age, and chemical treatment in zebrafish(48). See the supplemental methods for detailed parameters.

### ISH

Brain samples were sectioned on a cryostat at 16 μm onto four serial series. Tissue fixation parameters, probe synthesis, and ISH conditions were based on established protocols(49, 50). We used digoxigenin (DIG)-labeled probes for *Npas4a* and *Gabbr1a* genes. All individuals were processed simultaneously (one gene at a time) to avoid any potential colorimetric development differences across individuals due to batch effects. Riboprobes showed specific binding with high expression using the antisense probe, proportionally reduced expression in the 1:25 cold-competitor condition, and no expression in the sense and no probe conditions (Figure S1). See supplemental methods for detailed parameters.

### Brain Region Analysis

Brain section images were captured at 4X using a Nikon Eclipse monochrome camera (Qi2). For each brain region, we used Nikon NIS Elements Version 4.6 software to measure a standardized rectangular box within the borders of each brain region and measured the mean intensity of *npas4a* and *gabbr1a* expression within the box. The researcher (M.R.B.) was blinded to the treatment and strain conditions when collecting and analyzing images. We quantified gene expression by measuring optical density (OD) of the digoxigenin labeled probes, an established semi-quantitative measure of gene expression in other systems(49). For each slide, we normalized the mean intensity of all measures to the background (mean intensity of slide area not containing tissue), which produced a fractional transmittance value for each brain region in each section. Fractional transmittance was mathematically converted to optical density by the equation OD = 2-log(fractional transmittance). See supplemental methods for additional details.

### Statistics

All statistics were performed using SPSS software (Version 24). To analyze freezing behavior we used a repeated measures two-way ANOVA with strain and treatment group as between-subjects factors. For analyzing qRT-PCR gene expression we used a multivariate general linear model (GLM) with normalized *npas4a* and *gabbr1a* expression as dependent variables, and strain and treatment as between-subject factors. For analysis of ISH OD measurements we used a multivariate GLM with the OD of the five brain regions as dependent variables and strain and treatment group as between-subjects factors. There were not any effects of sex on learning and memory in a previous nor the current study (3-way repeated measures ANOVA: F_sex*trial_= 0.40 p = .531; F_sex_= 0.57 p = .456), so we removed sex as a variable to simplify the model(30). Individual groups were compared with simple effects testing. To account for multiple comparisons we applied the Benjamini-Hochberg correction to determine significance(51). For all significant differences (p < 0.05) we also report the effect sizes (Cohen’s d (d) for t-tests and partial eta-squared (ηp^2^) for ANOVAs (52). All effect sizes were medium or large effects(52–54).

## Results

### Contextual Fear Learning

In the conditioned fear response period during acquisition testing, there was a significant trial*treatment group interaction effect for freezing (*F*_1, 64_ = 54.86, *p* = 3.59*10^−10^, ηp^2^= .46). The alarm substance group showed increased freezing between trials at a faster rate than the DI control group (Figure 1). Additionally, there was a significant trial*strain*treatment group interaction (*F*_1, 64_ = 5.88, *p* = .018, ηp^2^= .08) where treated HSB fish increased freezing behavior at a faster rate than LSB fish. HSB fish exposed to alarm substance froze significantly more than LSB fish at trial two (*t*(32) = 4.23, *p* = 1.81*10^−4^, d = 1.45), but was not significant at trial one (*t*(32) = 1.05, *p* = .303). Full model results are presented in Table S2.

**Figure 1.**
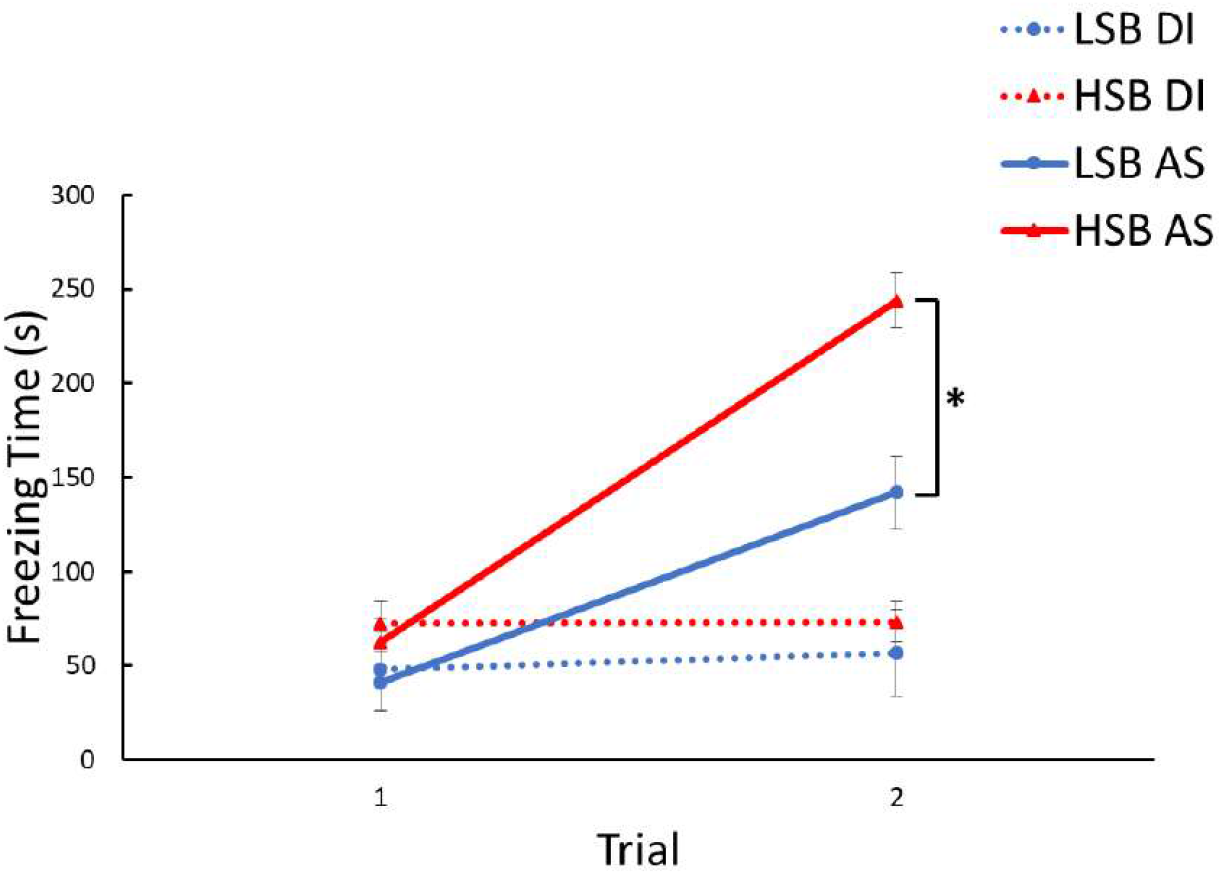
Acquisition of fear memory over two training trials. Freezing time was measured for high stationary behavior (HSB) and low stationary behavior (LSB) fish exposed to distilled water (DI) or alarm substance (AS). Points represent mean ± 1 standard error. * indicates *p* < .05 for within-treatment group comparison

### q*RT-PCR*

There was a significant effect of strain on both *npas4a* (*F*_1, 16_ = 11.72, *p* = .003, ηp^2^ = .42) and *gabbr1a* (*F*_1, 16_ = 7.29, *p* = .016, ηp^2^= .31) forebrain expression. There was a significant effect of treatment for *npas4a* (*F*_1, 16_ = 11.72, *p* = .003, ηp^2^= .42), but not *gabbr1a* (*F*_1, 16_ = 4.30, *p* = .055) expression. Full model results are presented in Table S3. In HSB fish, *npas4a* gene expression was significantly higher in the AS group compared to the DI group (*p* =.003, d = 2.34; Figure S2). There were no effects of treatment on *npas4a* expression in LSB fish (p=.918).

### In situ Hybridization

#### Treatment Effects on npas4a OD

There was a significant effect of treatment condition on *npas4a* OD in the Dm (*F*_2, 66_ = 6.20, *p* = .003, ηp^2^= .16), Dl (*F*_2, 66_ = 7.13, *p* = .002, ηp^2^= .18), Vv (*F*_2, 66_ = 3.38, *p* = .040, ηp^2^= .09), and Vs (*F*_2, 66_ = 3.93, *p* = .024, ηp^2^= .11). In the Dm, *npas4a* OD was significantly lower in DI water treatment group compared to both the baseline (*p* =.030, d = 0.67) and alarm substance group (*p* =.003, d = 1.04; Figure 2A). In the Dl, *npas4a* OD was significantly higher in the AS group compared to both the baseline (*p* =.042, d = 0.63) and DI water treatment group (*p* =.003, d = 1.05; Figure 2B). In the Vv, the AS group initially had a significantly higher OD compared to the baseline (*p* =.048, d = 0.59) and DI groups (*p* =.018, d = 0.71), however this was not significant after BH correction (*p* = .072, .054 respectively; Figure S3). In the Vs, *npas4a* OD was significantly lower in the DI group compared to both the baseline (*p* =.039, d =0.62) and AS treatment group (*p* =.033, d = 0.74; Figure 2C). In the Vd, *npas4a* OD was significantly higher in the AS group compared to the DI group for LSB fish only (*p* =.002, d = 1.00; Figure S3).

**Figure 2.**
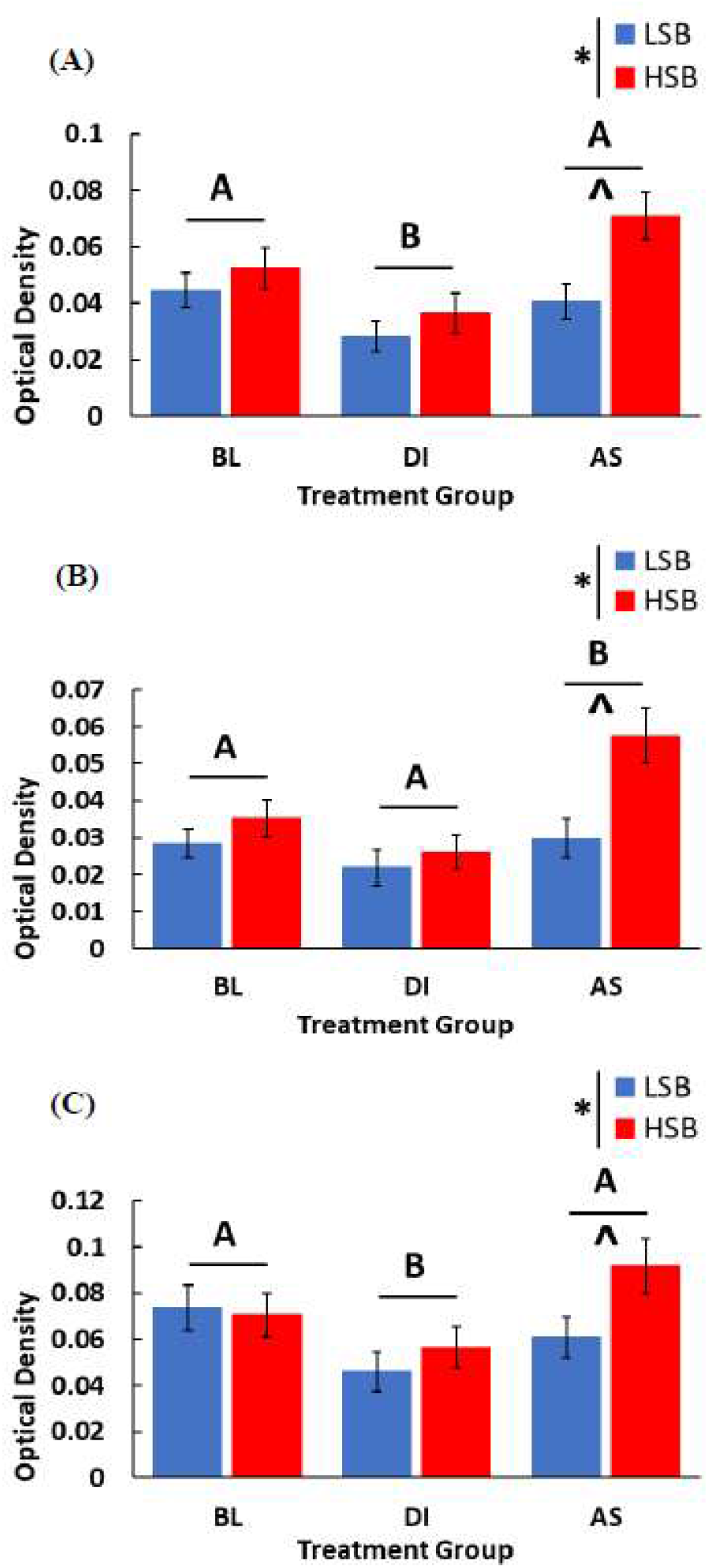
Expression of *npas4a* in the Dm (A), Dl (B), and Vs (C). We measured expression of high stationary behavior (HSB) and low stationary behavior (LSB) fish at baseline (BL) or exposed to either alarm substance (AS) or distilled water (DI) during training. Bars represent mean ± 1 SE. Bars labeled with different letters indicate p < .05. * indicates a significant strain main effect. ^ indicates a significant within-treatment group strain difference.

#### Strain Effects on Npas4a OD

There was a significant main effect of strain on the OD of *npas4a* in the Dm (*F*_1, 66_ = 7.66, *p* = .007, ηp^2^= .10), Dl (*F*_1, 66_ = 8.82, *p* = .004, ηp^2^= .12), and Vv (*F*_1, 66_ = 5.16, *p* = .026, ηp^2^= .07). HSB fish overall had higher OD of *npas4a* in each of the three brain regions. Additionally, HSB fish exposed to AS had significantly higher *npas4a* OD compared to LSB fish exposed to AS in the Dm (*p* =.001, d = 1.25), Dl (*p* =.001, d = 1.65), and Vs (*p* =.039, d = 0.65; Figures 2A-C). Full model results are presented in Table S4.

#### Strain Specific Treatment Effects on gabbr1a OD

For *gabbr1a* OD, there were significant strain*treatment group interaction effects in the Dm (*F*_1, 66_ = 3.31, *p* = .043, ηp^2^= .09), Vv (*F*_1, 66_ = 7.70, *p* = .001, ηp^2^= .19), Vd (*F*_1, 66_ = 6.95, *p* = .002, ηp^2^= .17), and Vs (*F*_1, 66_ = 3.89, *p* = .025, ηp^2^= .11). For each of those regions, there were no significant differences between any treatment groups for HSB fish. However, for LSB fish the DI group had significantly lower *gabbr1a* OD compared to the BL (*p* =.003, d = 1.37) and AS (*p* =.024, d = 1.00) groups in the Dm, BL (*p* =.023, d = 1.02) and AS (*p* =.003, d = 1.60) groups in the Vv, and the BL (*p* =.015, d = 1.06) and AS (*p* =.003, d = 1.37) groups in the Vd (Figure S4). The BL group had a significantly higher *gabbr1a* OD compared to the DI (*p* =.003, d = 1.71) and AS (*p* =.030, d = 0.99) groups in the Vs. Full model results are presented in Table S5.

## Discussion

Expression of neural plasticity-related genes (e.g. *npas4, gabbr1a*) has been broadly implicated as a key process underlying alternative stress coping styles, but has not been investigated related to proactive-reactive differences in learning and memory (20–22, 26, 27, 55, 56). Consistent with previous findings, we found that reactive (HSB) zebrafish showed an increased conditioned fear response relative to proactive (LSB) individuals (Figure 1)(30). Further, we found that *npas4a* expression was significantly higher in several key forebrain regions of reactive zebrafish. Altogether, our findings suggest that *npas4a* plays a similar role in learning and memory as its mammalian homolog, and may be an important regulator of proactive-reactive differences in learning and memory.

ISH analysis showed that *npas4a* expression was significantly higher in reactive fish in the Dm, Dl, and Vs (Figures 2A-C). The Dm (BLA), Dl (HIP), and Vs (BNST) are key sites of experience-dependent plasticity and integral to fear learning and memory across species(32–35). Similar to rodents, lesioning the teleost Dm and Dl impairs the formation of new fear and contextual memories (33, 57–59). Our findings suggest that *npas4a*-dependent plasticity within these brain regions may be a key underlying mechanism regulating differences in fear learning and memory capabilities between stress coping styles. In a prior study using the same conditioning paradigm, we showed that reactive zebrafish acquired a conditioned fear response faster than proactive zebrafish (cite). The higher activity-dependent expression of *npas4a* in reactive individuals observed in this study may promote higher levels of neural plasticity, resulting in salient and fearful experiences to be encoded into memory more quickly(28, 60). We predict that *npas4a* knockout experiments would produce similar learning and memory deficits as in rodents, and are needed to establish a direct causal role in zebrafish. More recently, specific glutamatergic populations of Dm cells have been shown to be required for fear conditioning (32). Our study is not able to distinguish between cell types expressing *npas4a* and would be needed to better characterize the specific circuits regulating proactive-reactive differences in learning. In selectively bred proactive and reactive trout, these telencephalic forebrain regions have also been shown to display differing monoaminergic and cortisol responses to acute stress(61, 62). This suggests that higher expression of *npas4a* in these brain regions may play important roles in constraining variation across a number of behavioral contexts.

While the BNST has been shown to be important for aversive learning in rodents(63, 64), the function of the Vs and specifically of *npas4a* expression in the Vs is not well understood in regards to learning and memory. We found that similar to the Dm and Dl, *npas4a* expression within the Vs is likely important for fear learning, and is associated with differences between proactive and reactive stress coping styles. Supporting this, a previous study found that increased activity and *npas4* expression in a population of corticotropin-releasing factor neurons in the BNST was associated with increased stress resiliency and prevention of a post-traumatic stress disorder-like phenotype in rodents(65). This suggests that *npas4a* expression in the Vs may play an important role in how individuals experience and cope with stress differently. Interestingly, the Vs has been shown to have connections with both the Dm and Dl, and to the hypothalamus and other brainstem areas that are essential for eliciting behavioral and endocrine stress responses. While this study only assessed gene expression across select forebrain structures, future studies should investigate other downstream structures and consider the role of glucocorticoids and the hypothalamus-pituitary-adrenal axis (hypothalamus-pituitary-interrenal in teleosts). This is particularly promising as glucocorticoid differences have been well-characterized between proactive and reactive stress coping styles(3, 66–68), though to a lesser extent related to learning and memory.

The DI treatment groups showed significantly lower *npas4a* expression compared to the AS treatment group in the Dm, Dl, and Vs (Figure 2a, 2c). This suggests *npas4*a is expressed in a treatment-specific manner associated with the learned conditioned fear response in the AS group. Unexpectedly, *npas4a* expression in the DI group was significantly lower than the BL group in the Dm and Vs. Other studies have found that acute injection of corticosterone or chronic restraint and social isolation stressors can decrease *npas4* expression in the rodent prefrontal cortex and hippocampus and lead to a variety of behavioral deficits including learning and memory(69–71). It is unclear whether this decrease in expression is maladaptive, or whether it is an adaptive homeostatic response to stress(72). It is unlikely that our results can be explained by physical isolation, as the baseline group was also socially isolated for the same duration. However, it is possible that handling stress could explain the reduction in *npas4a* expression for the DI group.

While qRT-PCR findings showed strain effects in *gabbr1a* expression, there were no strain differences in any of the analyzed brain regions for the ISH analysis. This suggests that the strain differences in forebrain *gabbr1a* expression are driven by other brain regions not investigated in this study. Therefore, *gabbr1a* expression within the Dm, Dl, Vv, Vs, and Vd does not appear to be associated with development of a conditioned fear response. Other studies have suggested that GABAergic signaling may be more important for consolidation, reconsolidation, or extinction of fear memories(73). Future studies should assess how GABA B receptor expression may influence other phases of fear conditioning, or other paradigms using positive reinforcement.

Learning to predict and cope with potentially dangerous environments is essential to an individual’s survival. Proactive and reactive stress coping styles represent alternative strategies for coping with stress and differ in a number of behavioral contexts, including learning and memory. Our study suggests that brain-region specific expression patterns of *npas4a* may underlie differences in fear learning between proactive and reactive stress coping styles. These findings advance our understanding of the neuromolecular mechanisms underlying stress-coping style differences in cognition and highlight neuroplasticity’s key role in regulating alternative adaptive behavioral responses to stress. Additionally, as proactive and reactive individuals share potentially conserved mechanisms underlying other stress coping behaviors, this suggests that these brain regions may also constrain behavioral variation in a number of disparate contexts.

## Acknowledgements

We are grateful to D. Revers, S. Roundtree, A. Parks, and N. Mohamed for zebrafish husbandry. Thank you to A. Goodman for assisting in collecting samples and K. Rushlau for assisting with molecular experiments. We are grateful to members of the Wong lab for comments on a prior version of this manuscript.

## Declarations

- The manuscript has been reviewed and approved by all listed authors for publication.
- All procedures were approved by the Institutional Animal Care and Use Committee of University of Nebraska at Omaha/University of Nebraska Medical Center (17-070-00-FC, 17-064-08-FC).
- This study was supported by funds from NSF (IOS-1942202), Nebraska EPSCoR First Award (OIA-1557417), Nebraska Research Initiative, and University of Nebraska Omaha (UNO) start-up grants to RYW. Funds were also provided by the UNO Biology Department, Rhoden Summer Graduate Fellowship, and the Graduate Research and Creative Activities Grants to MRB.
- The authors declare no competing interests
- MRB and RYW conceived and designed the experiments, and wrote the manuscript. MRB collected and analyzed the data.

## Supplementary Information

### Methods

#### qRT-PCR

We homogenized the tissue in Tri reagent (Sigma) and zirconium oxide beads in a Bullet Blender (NextAdvance) and extracted the RNA through column filtration (RNeasy Micro Plus Kit, Qiagen). RNA was subsequently converted to cDNA (Superscript IV First-Strand Synthesis System, Invitrogen) and purified (Millipore Amicon Ultra −0.5 mL 30 K Centrifugal Filters Devices). We ran qRT-PCR reactions on a QuantStudio 7 Flex Real-Time PCR system (Applied Biosystems) using PowerUp SYBR Green Master Mix (Applied Biosystems). A 131 base pair *npas4a* amplicon was created using 5’-CACCTCGGACACTCAATGGT-3’ (F) and 5’-AACAAGCGATCTGTGTCAGGT-3’ (R) as primers. A 198 base pair *gabbr1a* amplicon was created using 5’-CCCAGAGACGGAGGGATACG-3’ (F) and 5’-CGGGCACATCATCAAGCATCT-3’ (R) as primers. The parameters for both genes were as follows: 2 minutes at 50°C, 2 minutes at 95°C, followed by 40 cycles of 15 seconds of 95°C and 1 minute of 60°C. Primer concentration was 5 pmole/μl for both genes.

#### Tissue Section Processing

All series were simultaneously post-fixed in cold 4% paraformaldehyde/PBS solution, washed in PBS and acetylated in 0.25% acetic anhydride/triethanolamine. Then, slides were washed in 2X standard saline citrate, dehydrated in increasing ethanol series and stored at −80 °C.

#### Probe Synthesis

To quantify *npas4a* and *gabbr1a* we used digoxigenin (DIG)-labeled RNA probes. A 402 base pair *npas4a* DIG probe template was subcloned by using primer pair 5’-TTCTGTAGCGTCCAATCGGC-3’and 5’-ACTTCCACTCCCATCTTTGCG-3’. The 390 base pair *gabbr1a* probe template was subcloned by using primer pair 5’-AAGGATGAGCGCAATGTAGA-3’and 5’-CTGTTCCTGAGTCAGTCCTC-3’. Riboprobes were generated using a 1:3 ratio of UTP and DIG-UTP (Roche). After probe synthesis, we removed unincorporated nucleotides via column filtration according to manufacturer’s protocol (Megaclear, Ambion).

#### In situ Hybridization

Slides were prehybridized with a solution containing 50% formamide, 5X SSC, 5X Denhardt’s solution, 250 μg/ml yeast tRNA, and 500 μg/ml herring sperm DNA for 5 hours at 60°C in a hybridization chamber containing chamber buffer solution (50% formamide, 2X SSC). Then we hybridized the slides overnight at 67°C with fresh prehybridization solution containing 340 ng of *npas4a* antisense or 380 ng of *gabbr1a* riboprobe per slide. Following hybridization we performed two washes in 2X SSC at room temperature for *npas4a* (one wash in 2X SSC at 60°C, one wash in 2X SSC at room temperature for *gabbr1a*), then RNase A treated the slides (0.5M NaCl, 10 mM Tris pH 8.0, 2.25 mM EDTA, 0.2 μg/ml RNase A), followed by increasingly stringent washes (2X, 1X, 0.5X, 0.25X SSC) and then a final wash in Buffer B1 (100 mM Tris pH 7.5, 150 mM NaCl). Sections were then incubated overnight at 4°C with Anti-Digoxigenin AP antibody (Roche). After antibody incubation we washed sections twice in Buffer B1 and then blocked endogenous alkaline phosphatase activity with a 30 minute wash in Buffer B3 (100mM Tris pH 9.5, 100 mM NaCl, 50 mM MgCl2, 5 mM levamisole) in the dark. We used colorimetric detection using NBT/BCIP stock solution (Roche). The colorimetric reaction was stopped (80 minutes for *Npas4a* and 12 hours for *Gabbr1a*) by rinsing sections three times in ultrapure type 1 water and then progressively dehydrating sections in ethanol (25%, 50%, 70%, 95%).

#### Brain Region Analysis

The light settings were set to the maximum, and two 1/16 filters were placed over the light source to keep consistency across days. The measuring box was always placed in the middle of the brain region on the dorsal-ventral plane, excluding the midline. We measured the mean intensity bilaterally if available, and averaged all of the intensities for each individual for each brain region. Depending on the size of the brain region, the number of sections averaged per individual ranged from two to six consecutive sections. Consecutive sections were 48 μm apart. The anterior commissure was identified as a landmark for each of the brain regions. We measured the Dm (13003.92 μm^2^) and Dl (13003.92 μm^2^) for 1-2 sections prior to and 3-4 sections following the anterior commissure. We measured the Vv (9907.28 μm^2^) and Vd (9907.28 μm^2^) for 3-4 sections preceding the anterior commissure. We measured the Vs (9907.28 μm^2^) for the slice containing the anterior commissure and 1-2 following it.

---

**Table S1.** Brain region terminology, abbreviations, and putative tetrapod homologue regions.

| Teleost Region | Abbreviation | Putative Tetrapod Homologue |
| --- | --- | --- |
| Area dorsomedialis telencephali | Dm | Basolateral amygdala |
| Area dorsolateralis telencephali | DI | Pallial hippocampus |
| Area ventroventralis telencephali | Vv | Lateral septum |
| Area dorsoventralis telencephali | Vd | Striatum |
| Ventralis supracommissuralis telencephali | Vs | Bed nucleus of the stria terminalis |

**Table S2.** Results of repeated measures GLM for the acquisition learning phase for freezing time.

| Freezing Time |  |
| --- | --- |
|  | F<sub>(p, ηp<sup>2</sup>)</sub> |
| Within-Subjects Effects (df = 1, 64) |  |
| Trial | **62.82** (4.36*10<sup>-11</sup>, .50) |
| Trial*Strain | 3.89 (.053) |
| Trial*Treatment | **54.86** (3.59*10<sup>-10</sup>, .46) |
| Trial*Strain*Treatment | **5.88** (.018, .08) |
| Between Subjects Effects (df = 1, 64) |  |
| Intercept | **179.53** (3.08*10<sup>-20</sup>, .74) |
| Strain | **8.92** (.004, .12) |
| Treatment | **18.78** (5.30*10<sup>-5</sup>, .23) |
| Strain*Treatment | 2.18 (.144) |

Bold text indicates $p < 0.05$

**Table S3.** Results of multivariate GLM for forebrain expression of *npas4a* and *gabbr1a* from qPCR.

|  | *npas4a* | *gabbr1a* |
| --- | --- | --- |
|  | F<sub>(p, ηp<sup>2</sup>)</sub> | F<sub>(p, ηp<sup>2</sup>)</sub> |
| Intercept | **393.93** (1.08*10<sup>-12</sup>, .96) | **364.98** (1.94*10<sup>-12</sup>, .96) |
| Strain | **11.72** (.003, .42) | **7.29** (.016, .31) |
| Treatment | **11.72** (.003, .42) | 4.30 (.055) |
| Strain*Treatment | 2.32 (.147) | 3.88 (.066) |

Bold text indicates $p < 0.05$

**Table S4.**
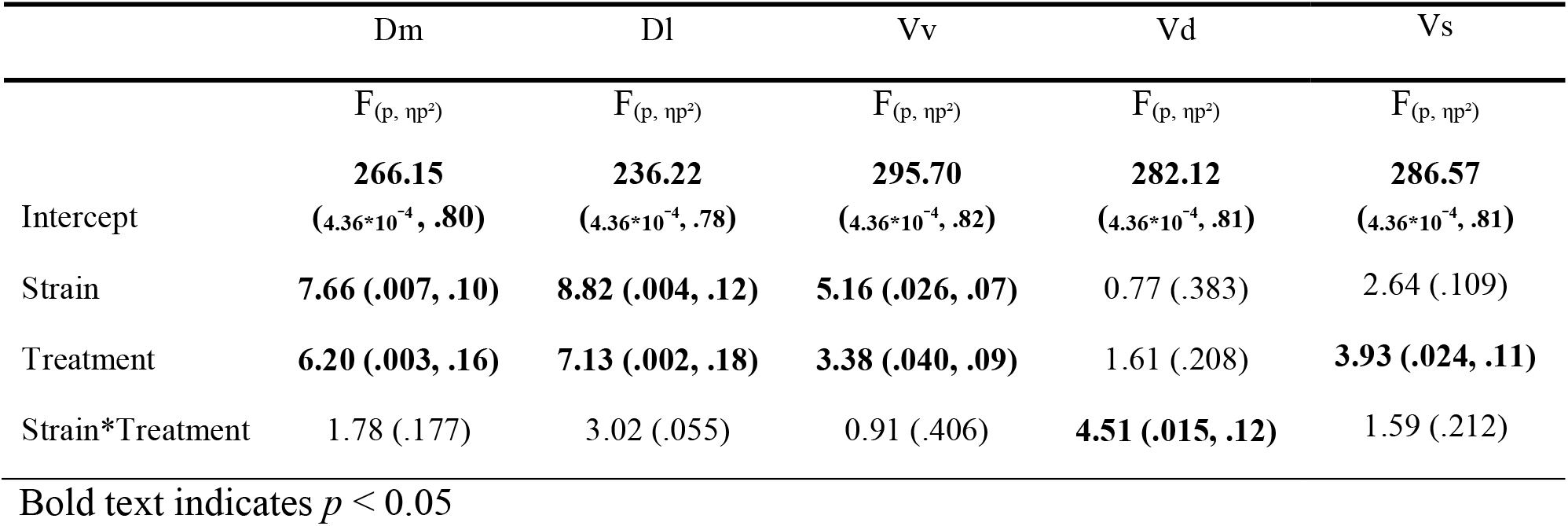
Results of multivariate GLM of *npas4a* optical density across the five forebrain regions.

**Table S5.**
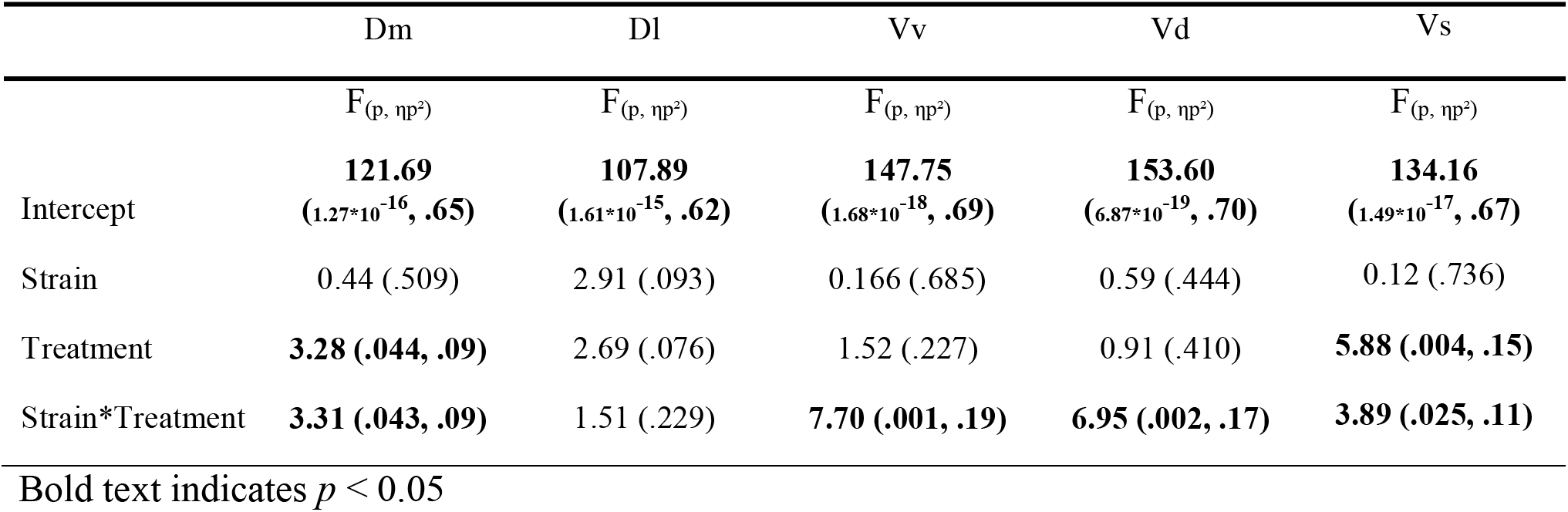
Results of multivariate GLM of gabbr1a optical density across the five forebrain regions.

|  | Dm | DI | Vv | Vd | Vs |
| --- | --- | --- | --- | --- | --- |
| | $F_{(p, \eta p^2)}$ | $F_{(p, \eta p^2)}$ | $F_{(p, \eta p^2)}$ | $F_{(p, \eta p^2)}$ | $F_{(p, \eta p^2)}$ |
| Intercept | **121.69**<br>($1.27 \times 10^{-16}$, .65) | **107.89**<br>($1.61 \times 10^{-15}$, .62) | **147.75**<br>($1.68 \times 10^{-18}$, .69) | **153.60**<br>($6.87 \times 10^{-19}$, .70) | **134.16**<br>($1.49 \times 10^{-17}$, .67) |
| Strain | 0.44 (.509) | 2.91 (.093) | 0.166 (.685) | 0.59 (.444) | 0.12 (.736) |
| Treatment | **3.28 (.044, .09)** | 2.69 (.076) | 1.52 (.227) | 0.91 (.410) | **5.88 (.004, .15)** |
| Strain*Treatment | **3.31 (.043, .09)** | 1.51 (.229) | **7.70 (.001, .19)** | **6.95 (.002, .17)** | **3.89 (.025, .11)** |

Bold text indicates $p < 0.05$

**Figure S1.**
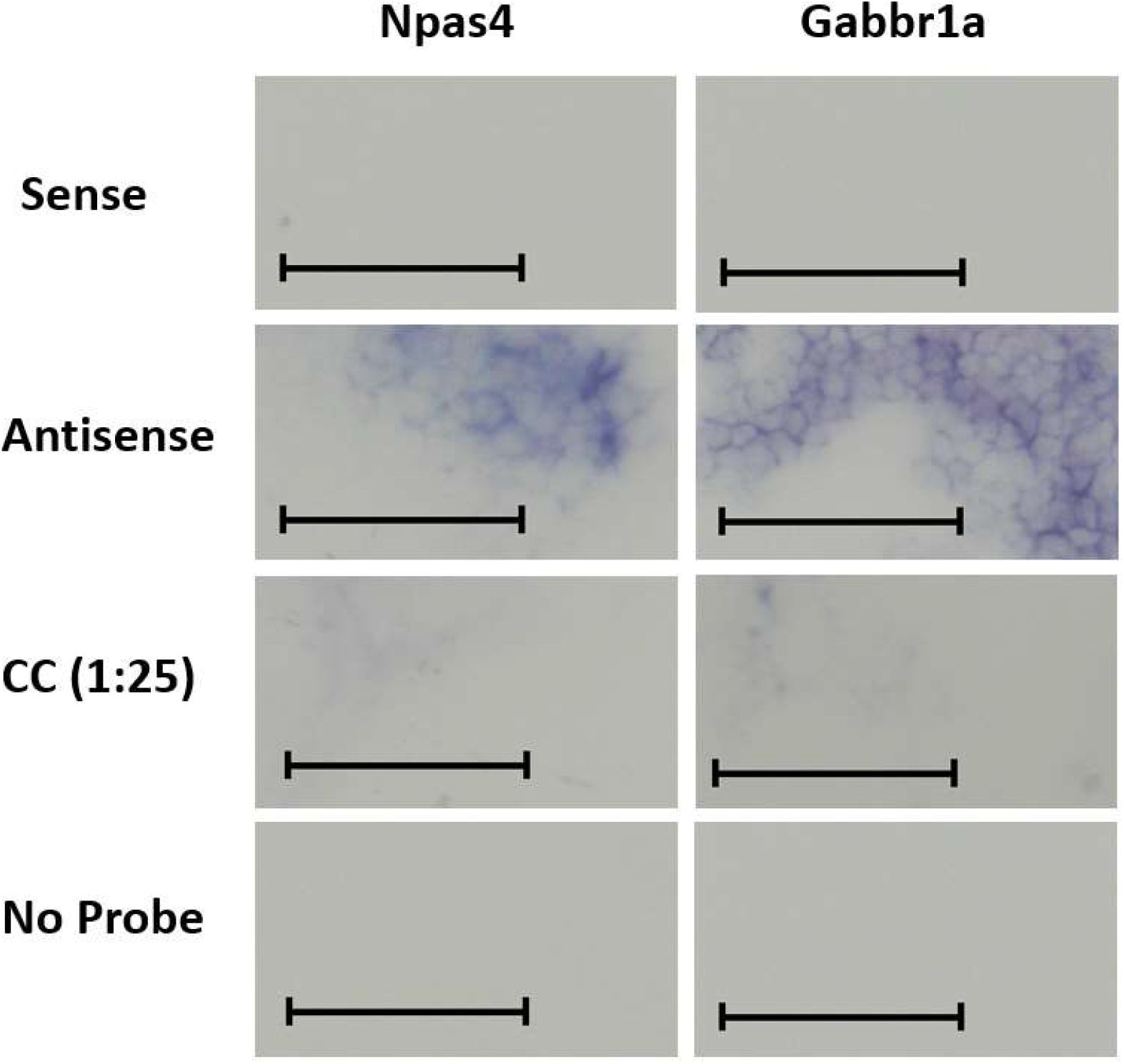
Pilot *in situ* hybridization results for the *Npas4a* and *Gabbr1a* genes. There was strong signal in the antisense, proportionally reduced signal in the cold-competitor (1:25 ratio of DIG-labeled to unlabeled riboprobe), and negligible signal in the sense and no probe permutations. Scale bars represent 50 um.

**Figure S2.**
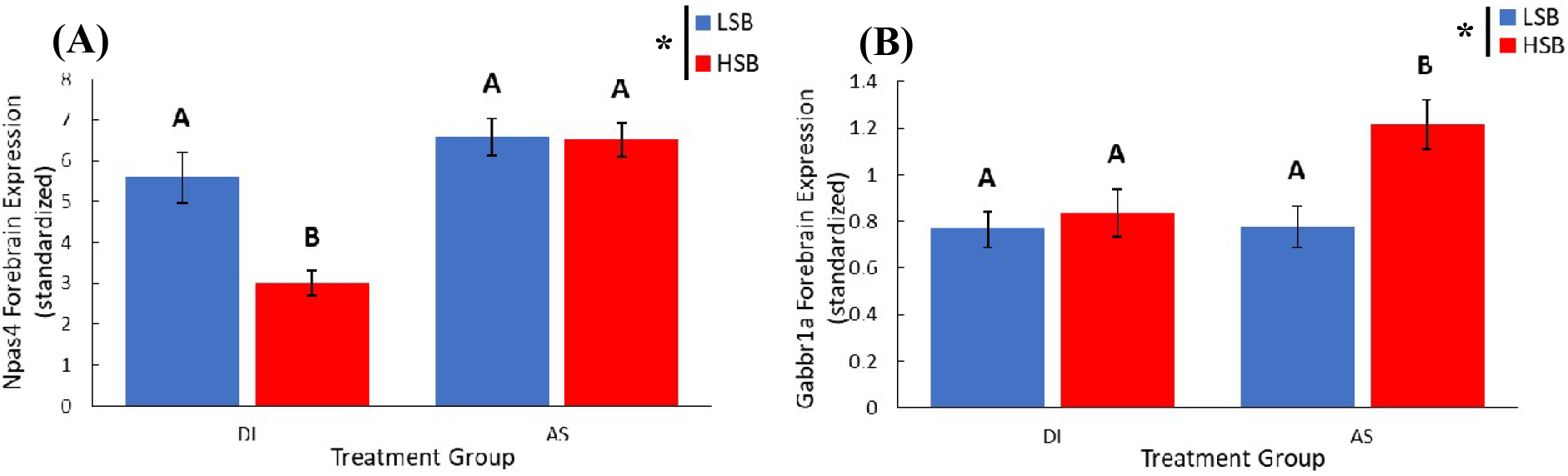
*npas4a* (A) and *gabbr1a* (B) forebrain expression standardized to *ef1a*. We measured expression of high stationary behavior (HSB) and low stationary behavior (LSB) fish that were exposed to either alarm substance (AS) or distilled water (DI) during training. Bars represent mean ± 1 SE. Bars labeled with different letters indicate p < .05. * indicates a significant strain main effect.

**Figure S3.**
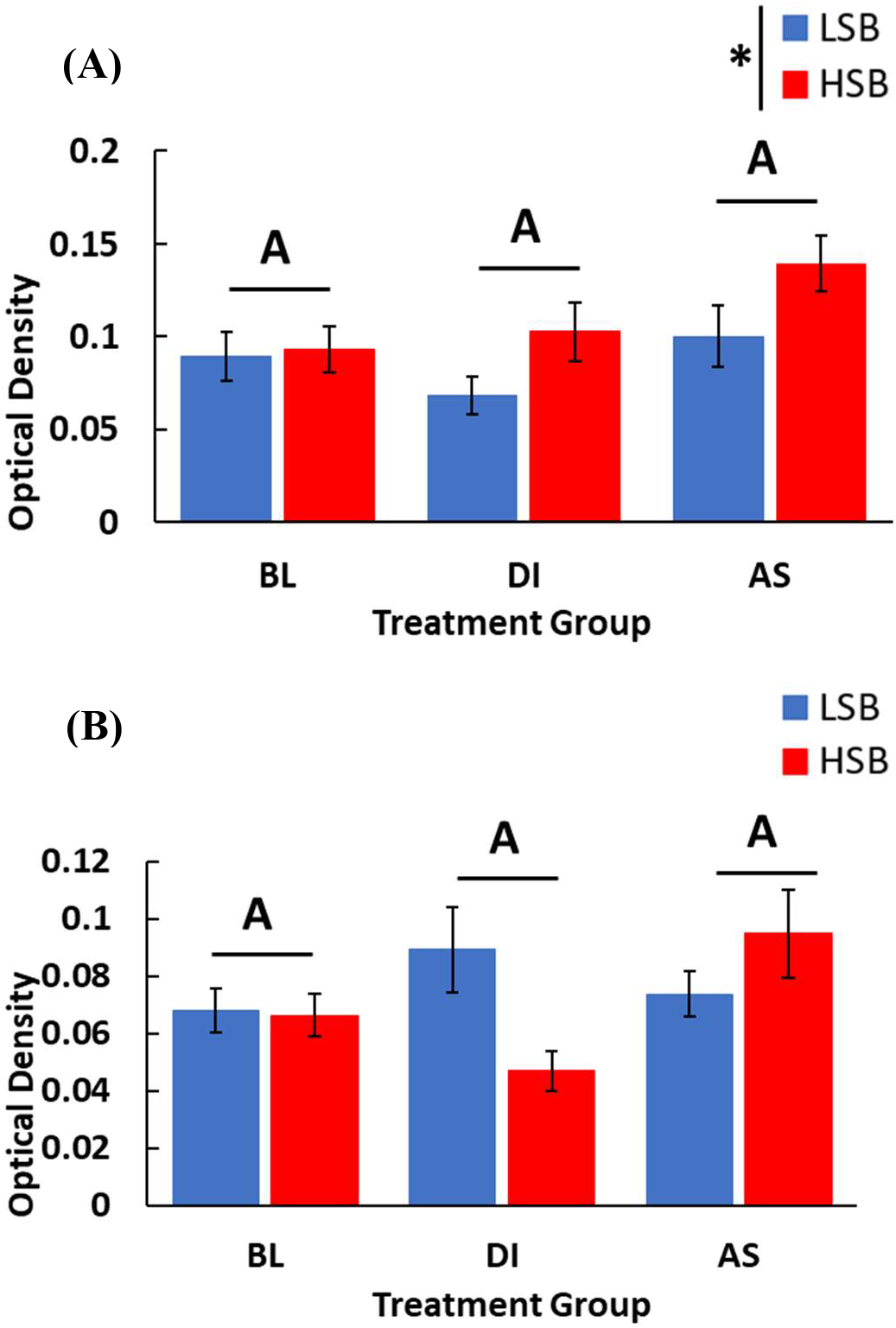
Expression of *npas4a* in the Vv (A) and Vd (B). We measured expression of high stationary behavior (HSB) and low stationary behavior (LSB) fish at baseline (BL) or exposed to either alarm substance (AS) or distilled water (DI) during training. Bars represent mean ± 1 SE. Bars labeled with different letters indicate p < .05. * indicates a significant strain main effect.

**Figure S4.**
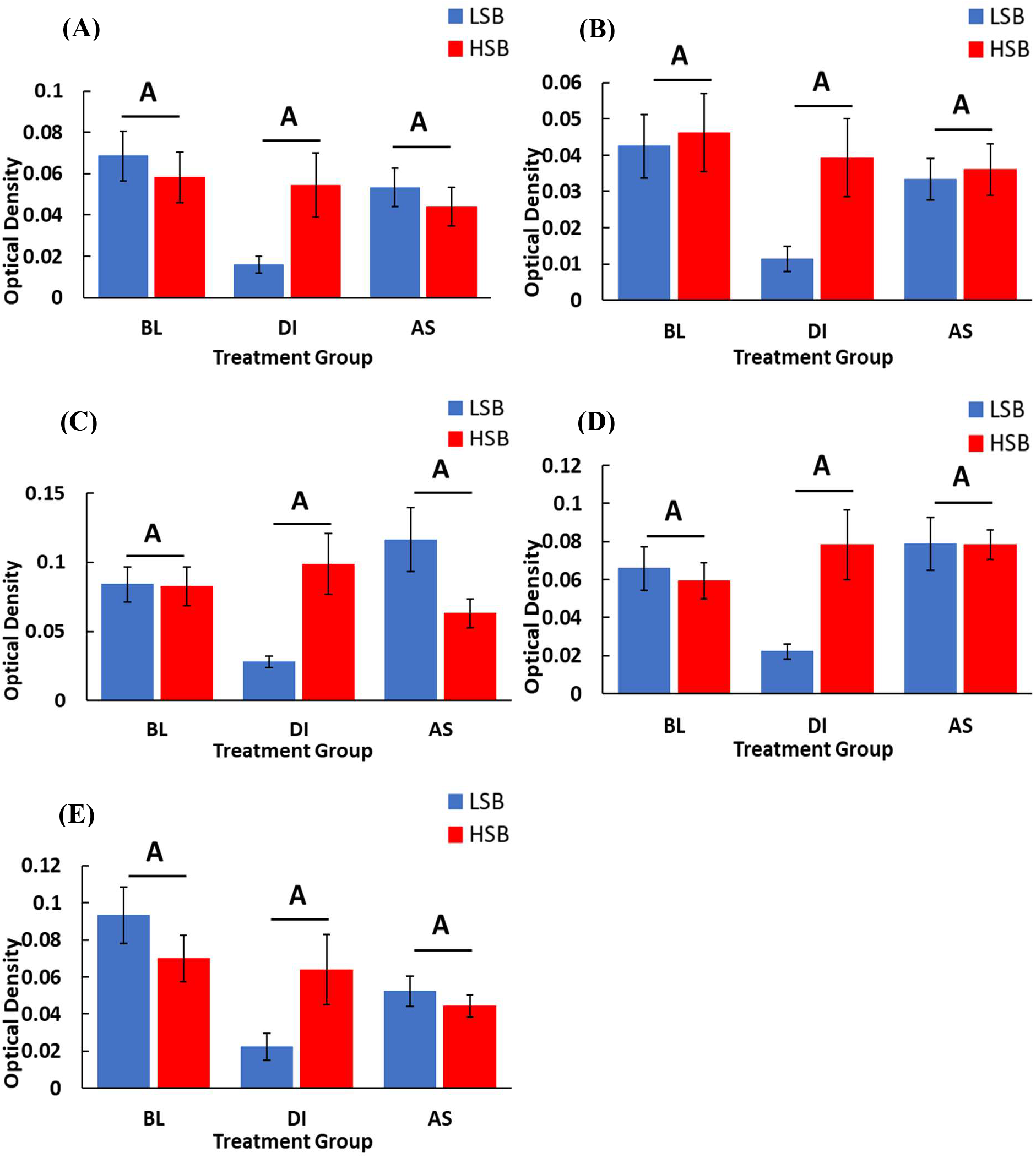
Expression of *gabbr1a* in the Dm (A), Dl (B), Vv (C), Vd (D), Vs (E). We measured expression of high stationary behavior (HSB; B) and low stationary behavior (LSB; A) fish at baseline (BL) or exposed to either alarm substance (AS) or distilled water (DI) during training. Bars represent mean ± 1 SE. Bars labeled with different letters indicate p < .05. When split by strain, LSB fish exposed to DI water had significantly lower *gabbr1a* OD compared to the baseline and AS groups. There were no treatment group differences in the HSB group.

